# Universal preprocessing of single-cell genomics data

**DOI:** 10.1101/2023.09.14.543267

**Authors:** A. Sina Booeshaghi, Delaney K. Sullivan, Lior Pachter

**Affiliations:** Division of Biology and Biological Engineering, California Institute of Technology, Pasadena, CA, USA; UCLA-Caltech Medical Scientist Training Program, David Geffen School of Medicine, University of California, Los Angeles, Los Angeles, CA, USA; Department of Computing & Mathematical Sciences, California Institute of Technology, Pasadena, CA, USA

## Abstract

We describe a workflow for preprocessing a wide variety of single-cell genomics data types. The approach is based on parsing of machine-readable *seqspec* assay specifications to customize inputs for *kb-python*, which uses *kallisto* and *bustools* to catalog reads, error correct barcodes, and count reads. The universal preprocessing method is implemented in the Python package *cellatlas* that is available for download at: https://github.com/cellatlas/cellatlas/.

## Introduction

A frequent refrain in single-cell genomics is that “uniform processing” of data is desirable and important. However when contrasted against the large number of single-cell genomics assays (Klein et al. 2015; Hagemann-Jensen et al. 2020; Zheng et al. 2017; Rosenberg et al. 2018; W. Xu et al. 2022; Gehring et al. 2020; Stoeckius et al. 2017; Schraivogel et al. 2020), and given that ideally analysis methods should be optimized and tailored for the assays to which they will be applied, the question arises why is uniform processing desirable? For example, in the case of single-cell RNA-seq it may be interesting to study cell-type isoform specificity (Booeshaghi et al. 2021), whereas for single-cell ATAC-seq the notion of isoforms is meaningless because the assay is based on genomic DNA, not cDNA, sequencing (Cusanovich et al. 2015). The reason uniform preprocessing can be important, is that with increasing amounts of data being generated (Svensson, da Veiga Beltrame, and Pachter 2020), datasets are seldom analyzed in isolation, and to the extent possible, it is desirable to avoid batch effects resulting from assay- or data-specific methods. Thus, uniform processing is increasingly intended to mean uniform preprocessing^1^ of data so that meta-analyses can be shielded from batch effects (Zhang et al. 2021; Davis et al. 2018).

Uniform preprocessing of single-cell genomics data is possible because single-cell genomics protocols are all rooted in a handful of technologies. They can be organized according to their physical-isolation and molecular-capture modes: cellular contents can be isolated in wells (Picelli et al. 2014; Gierahn et al. 2017; Hagemann-Jensen et al. 2020), shells (Zheng et al. 2017; Macosko et al. 2015; Klein et al. 2015), or cells, where the cell (or nucleus) itself serves as a container (Rosenberg et al. 2018; Cao et al. 2017; Ma et al. 2020). Physical isolation, whether by wells, shells, or cells, is followed by wet-lab methods to capture molecular features. These features are tagged using barcodes that associate synthetic nucleotide sequences to individual objects such as cells, nuclei, pixels, or beads. These “molecular capture modes”, that may include RNA, DNA, proteins, or synthetic tags, are coupled with barcode sequences to produce libraries for sequencing.

The physical-isolation and molecular-capture methods used for an assay are reflected in the structure of reads produced by these methods (Figure 1). The *seqspec* assay specification (Booeshaghi, Chen, and Pachter 2023) provides a machine-readable format for describing this structure, thereby translating the organizational themes of single-cell genomics into a medium that can, in principle, facilitate automatic and universal preprocessing of single-cell genomics data. While several tools have been developed for preprocessing data from different single-cell RNA-seq assays (He et al. 2022; Battenberg et al. 2022; Melsted et al. 2021), we demonstrate that *seqspec* (Booeshaghi, Chen, and Pachter 2023), along with *kallisto bustools* (Melsted et al. 2021), *kITE* (Sina Booeshaghi et al. 2022) and *snATAK* (Sina Booeshaghi, Gao, and Pachter 2023), can in principle be used for preprocessing data from any single-cell genomics assay.

**Figure 1:**
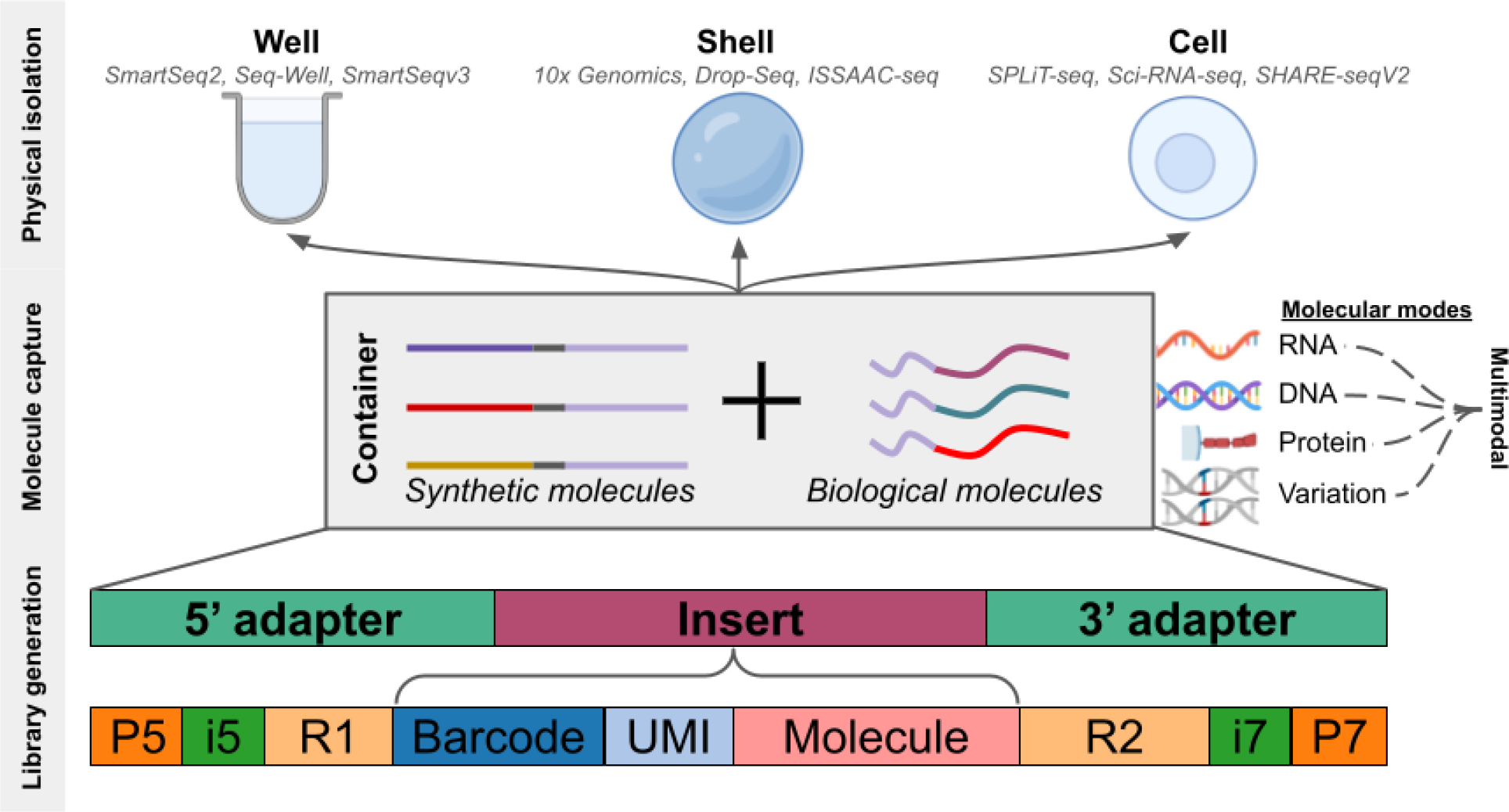
Single-cell genomics assays are variations on a theme of physical isolation, molecular capture, and library generation. The read structure shown here is provided as an example; read structure may vary depending on assay type.

## Results

To consistently preprocess single-cell genomics data, reads sequenced from libraries produced with different technologies must to be cataloged, error-corrected, and counted. Cataloging reads requires determining their sequence of origin by mapping them either to known or unknown categories. For example, single-cell RNA-seq reads might be aligned to a known set of transcripts to count RNA molecules. Similarly, in cases where the set of possible orthogonal barcodes is known *a priori*, as in PerturbSeq experiments (Schraivogel et al. 2020), cell assignment can be performed based on mapping to known barcode categories. On the other hand, unknown categories may be informed by the sequencing reads themselves, such as in ATAC-seq assays where accessible chromatin categories can be determined by the reads. The *kallisto bustools* workflow can build known and unknown categories for a wide range of assays such as RNA-seq, ATAC-seq, and synthetic and protein-tag based methods such as CITE-seq (Stoeckius et al. 2017), MultiSeq (McGinnis et al. 2019), and Clicktags (Gehring et al. 2020). Genomic and transcriptomic categories are generated for scRNAseq and synthetic tag libraries for orthogonal barcode methods while peak-categories are generated from the reads for chromatin accessibility libraries.

Universal preprocessing also requires identifying and handling sequence elements from reads in a consistent manner. Elements such as barcodes, unique molecular identifiers (UMIs), and genomic features such as cDNA must be accurately identified and appropriately parsed by preprocessing tools to ensure that cataloging, error correcting, and counting are correctly performed. To enable proper element identification and universal preprocessing, we use the *seqspec index* command (Booeshaghi, Chen, and Pachter 2023) to extract sequenced elements in a tool-specific manner and perform read cataloging against the previously-indexed categories. This indexing strategy correctly identifies and extracts relevant single-cell features in all assay types, a result of the consistent and controlled vocabulary of the *seqspec* specification.

Sequencing machines used to sequence the libraries produced with genomics assays may generate reads with errors in barcode sequences affecting the ability for barcode-barcode registration in multi-modal assays. Preprocessing tools that perform discordant barcode error correction strategies on each modality may lose barcodes due to mismatches in correction. To address this, universal preprocessing requires consistent error correction strategies. The *kallisto bustools* workflow achieves a good tradeoff between speed, memory, and barcode recovery for the Hamming-1 distance barcode error correction strategy and can be consistently applied to barcodes from multiple assays (Melsted et al. 2021).

After read cataloging and error correction, the final step in universal preprocessing is read counting. To quantify the number of features per cell in single-cell genomics assays, molecular counting approaches must be applied in a manner that balances accuracy, speed, and memory while being consistent between assays that do, and do not, have UMIs. We leverage the *kallisto bustools* counting approach as it performs UMI and read counting in a consistent way. This means that UMIs quantified in scRNAseq data are resolved in a manner that is consistent with reads quantification for scATAC-seq data (Sina Booeshaghi, Gao, and Pachter 2023).

Additionally, the expectation-maximization algorithm can be applied to disambiguate multi-mapping reads(Bray et al. 2016; Booeshaghi et al. 2021).

These universal principles guided our development of *cellatlas* for universal preprocessing of single-cell genomics data. The *cellatlas* program requires sequencing reads, genomic references, and a *seqspec* specification file. From there, *cellatlas* leverages *seqspec* functionality to auto generate the *kallisto* technology string that specifies the 0-index position of the cellular barcodes, molecular barcodes, and genomic features such as cDNA or genomic DNA. It also either points to, or generates, a barcode onlist compatible with the barcodes in the FASTQ files. Thus, *cellatlas* can be used to process data with a single command:

~~~
cellatlas build -o out -m rna -s spec.yaml -fb bcs.txt -fa genome.fa -g genome.gtf R1.fastq.gz R2.fastq.gz
~~~

To demonstrate the universality of *cellatlas* and its application to uniform preprocessing we applied *cellatlas* to preprocess nine datasets from three physical isolation methods which capture five different molecular modalities: spatial RNA, nuclear RNA, cytoplasmic RNA, accessible chromatin, synthetic tags, CRISPR guides, and cell surface proteins (see Methods).

Uniform preprocessing makes it straightforward to compare modalities within the same technology. Figure 2 shows a comparison of data from three different DOGMA-seq (Mimitou et al. 2021) modalities as assayed in (Z. Xu et al. 2022), all preprocessed with *cellatlas* and the tools it uses: *snATAK, kallisto bustools*, and *kITE*. Uniform preprocessing also allows for quantifying the efficiency of assaying the different modalities. The fact that the RNA and protein libraries have a similar number of UMIs captured for a given number of reads (Figure 2a,b) is consistent with similar dilutions of the libraries (1.99nM and 2.2nM respectively), whereas the tag library efficiency (Figure 2c) is distinctly different, likely as a result of dilution to 0.27nM. While the exact cause for the difference requires further investigation, the control for preprocessing that *cellatlas* provides points to an experimental cause for the difference. Uniform preprocessing also provides a direct comparison of the distinct modalities in terms of their detection efficiency (Figures 2d,e,f). The protein detection saturates because DOGMA-seq protein modality only assayed 163 proteins. The discrepancy between cells in which protein is detected but not ATAC peaks, versus those in which protein is detected but not RNA, is interesting and warrants further investigation.

**Figure 2:**
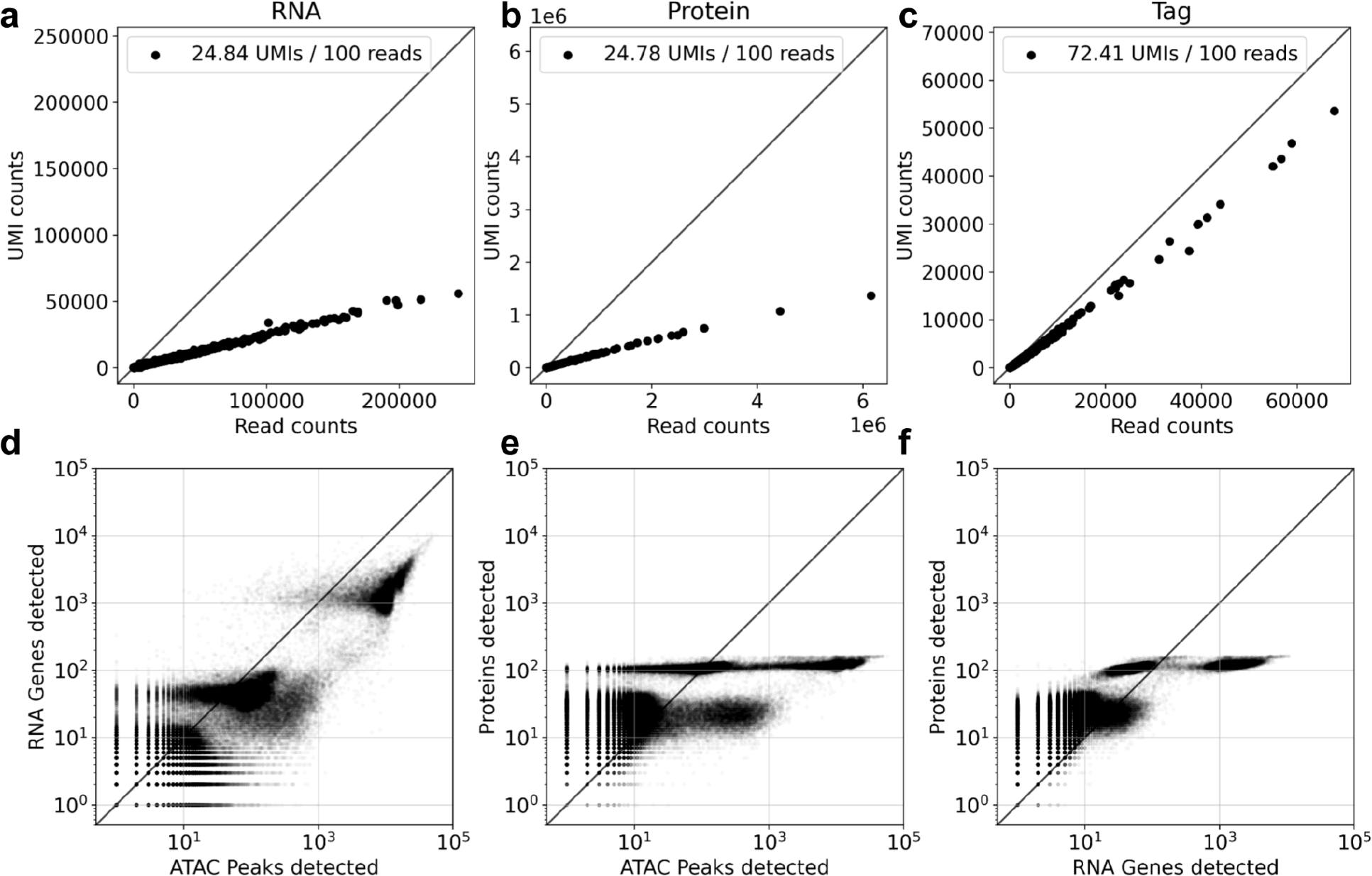
Efficiency of UMI detection as measured by relating the number of reads uniquely aligned for each cell to the number of UMIs for the a) RNA, b) protein, and c) tag modalities.

Uniform preprocessing also allows for comparing technologies. We compared single-cell RNA-seq and ATAC-seq produced with DOGMA-seq to those produced with 10x Multiome on PBMCs (see Methods). This cross-assay comparison was possible because we uniformly processed the data using the same read pseudoalignment algorithm, barcode error correction, and UMI/read counting. Figure 3 shows RNA-ATAC count statistics for DOGMA-seq alongside 10x Genomics Multiome for both datasets, and their constituent RNA and ATAC parts preprocessed using *cellatlas*. The DOGMA-seq assay appears to be more efficient than the 10x Multiome, and displays less background noise.

**Figure 3:**
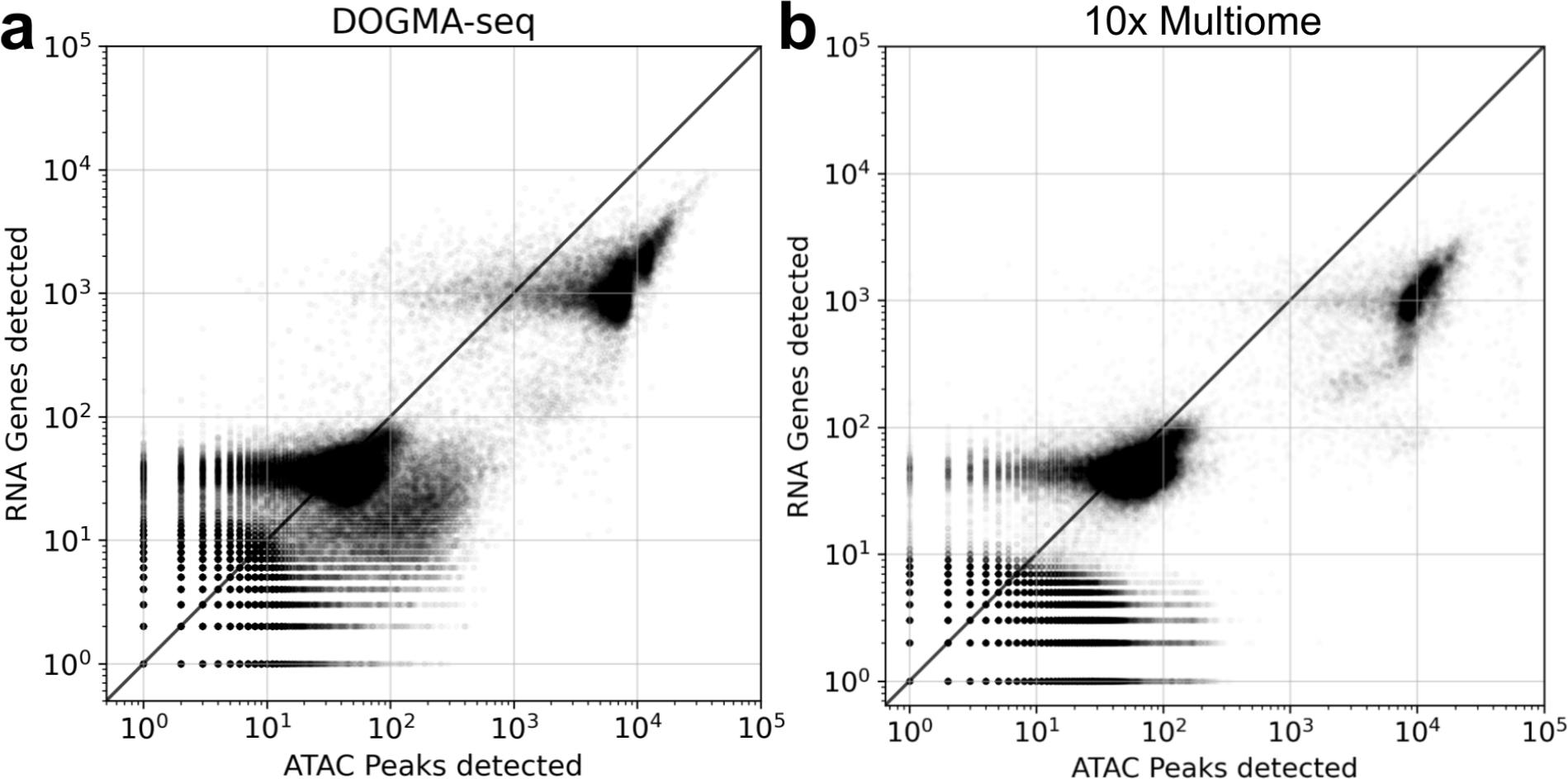
Cross-technology comparison of registered ATAC and RNA quantifications from PBMCs assayed with (a) DOGMA-seq and (b) 10x Multiome.

## Discussion

The importance of uniform preprocessing of single-cell genomics has been evident in the development of several workflows for multiple single-cell RNA-seq assays (Battenberg et al. 2022; He et al. 2022; Parekh et al. 2018; Tian et al. 2018). Here we have introduced an approach to uniform preprocessing that generalizes to many more assays and modalities by leveraging the modularity and flexibility of the *kallisto* and *bustools* programs, along with the *seqspec* standard for describing common elements in reads derived from single-cell genomics assays. Crucially, our approach enables quantitative comparisons both of distinct modalities within assays, as well as across technologies.

In addition to facilitating uniform analysis across many data types and studies, we anticipate that universal preprocessing methods will be useful in the development of new single-cell genomics assays, for example by helping to reveal cross-platform tradeoffs (Sina Booeshaghi, Gao, and Pachter 2023). Furthermore, uniform preprocessing will improve reproducibility of studies, because the association of a *seqspec* with an assay adds transparency to the protocol, and fast re-processing of data with *cellatlas* will allow users to quickly reproduce work that is currently a bioinformatics bottleneck for many projects. Finally, we envision that *cellatlas* will be useful for large-scale cell atlas coordination efforts.

## Acknowledgments

ASB and LP were partially funded by NIH 5UM1HG012077-02. DKS was partially funded by the UCLA-Caltech Medical Scientist Training Program (NIH NIGMS training grant T32 GM008042).

## Data & code availability

Example datasets that demonstrate the utility of *cellatlas* are available here: https://github.com/cellatlas/cellatlas/tree/main/examples. *cellatlas* is freely and openly available here: https://github.com/cellatlas/cellatlas/. All analysis notebooks can be found here: https://github.com/pachterlab/BSP_2023/

## Methods

### DOGMA-seq

DOGMA-seq FASTQ files were obtained from the Gene Expression Omnibus accession GSE200417. FTP links to SRA and FASTQ files were obtained with *ffq (Gálvez-Merchán et al. 2022)* by running *ffq –ftp GSE200417*. The DOGMA-seq ATAC and RNA modalities, for the 10x Multiome comparison in Figure 3, were subsampled to 400 million reads using the *seqkit* tool (Shen et al. 2016) *seqkit subsam*.

### DOGMA-seq preprocessing

All preprocessing was performed by running the *cellatlas* command with the FASTQ files, associated *seqspec*, and reference sequences. The *cellatlas* commands can be found within each example folder in the *cellatlas_info*.*json* file. RNA and DNA reads were mapped to using the GRCh38 genome and GRCh38-108 annotation.

### Efficiency calculation

In order to compute the sum of reads for each cell, we used *bustools count* with the *–cm* option. The resultant matrix is summed along the features to produce a vector for each cell. Then the sum of reads for each cell was plotted against the sum of UMIs for each cell. The slopes of the best fit lines were computed with the *sklearn* LinearRegression function.

## Footnotes

1 In the study of single-cell genomics data, the distinction between “preprocessing” and “processing” methods relates to the difference between those that operate on sequencing reads and those that operate on count matrices. The interchangeable use of these two terms has been a source of confusion for scientists and tool developers.

## Notes

https://github.com/cellatlas/cellatlas/

https://github.com/pachterlab/BSP_2023/

### Competing Interest Statement

The authors have declared no competing interest.

